# HologLev: Hybrid Magnetic Levitation Platform Integrated with Lensless Holographic Microscopy for Density-Based Cell Analysis

**DOI:** 10.1101/2020.12.10.420018

**Authors:** Kerem Delikoyun, Sena Yaman, Esra Yilmaz, Oyku Sarigil, Muge Anil-Inevi, Engin Ozcivici, H. Cumhur Tekin

## Abstract

In clinical practice, a variety of diagnostic applications require the identification of target cells. Density has been used as a physical marker to distinguish cell populations since metabolic activities could alter the cell densities. Magnetic levitation offers a great promise for separating cells at single cell level within heterogeneous populations with respect to cell densities. Traditional magnetic levitation platforms need bulky and precise optical microscopes to visualize levitated cells. Moreover, the evaluation process of cell densities is cumbersome, which also requires trained personnel for operation. In this work, we introduce a device (HologLev) as a fusion of magnetic levitation principle and lensless digital inline holographic microscopy (LDIHM). LDIHM provides ease of use by getting rid of bulky and expensive optics. By placing an imaging sensor just beneath the microcapillary channel without any lenses, recorded holograms are processed for determining cell densities through a fully automated digital image processing scheme. The device costs less than $100 and has a compact design that can fit into a pocket. We perform viability tests the device by levitating three different cell lines (MDA-MB-231, U937, D1 ORL UVA) and comparing them against their dead correspondents. We also tested the differentiation of mouse osteoblastic (7F2) cells by monitoring characteristic variations in their density. Lastly, MDA-MB-231 cells exposed to a chemotherapy drug are separated from original cell lines in our platform. HologLev provides cost-effective, label-free, fully automated cell analysis in a compact design which could be highly desirable for laboratory and point-of-care testing applications.

## INTRODUCTION

Global healthcare becomes more decentralized each year owing to the cost of delivering effective diagnostics and therapeutic services at central medical institutions which gradually lead to adapt some of the most crucial diagnostic tests into portable, rapid and economically desirable devices^1^. However, the identification of target cells within heterogeneous cell populations or bodily fluids (e.g., blood) remains as important a goal as the same for a variety of clinical testing. Thus, gold standard techniques for cell separations heavily rely on some well-known biophysical or biochemical aspects of cells, such as size, density, or surface receptor markers of cells^2^. Biochemical or immunoassays require high-cost reagents, bulky and precise instrumentation and trained personnel, conditions which can be supported at institutional level. Furthermore, cell labeling protocols are labor intensive and expensive procedures^3^. On the other hand, separating target cells using biophysical properties is a low-cost, rapid, and simple method to be deployed in a wide range of facilities. However, some of the biophysical properties such as size difference may not be suitable for purifying target cells ^4^. One of the most widely used techniques for cell separation is the differences in the density^5^. Density differences within cell populations may occur based on biochemical and genetic factors, such as differential gene expression and energy consumption ^6, 7^. Although such differences in density among cell populations are quite low, they can be utilized for separation at single cell level ^8^. Most widely used methods for determining density of cell populations are Ficoll gradient centrifugation^9^ and dielectrophoretic field-flow fractionation ^10^, none of which allow measurement of cell density at single cell level thus, it is unlikely to separate target organism at single cell level by using these techniques. Recently, magnetic levitation has shown superior performances over traditional density-based separation techniques that it enables measurement of cell density at single cell level by precisely balancing forces acting on each cell at micro scale^8^. Each cell is levitated to suspend at a unique height directly related with the density. Thus, magnetic levitation offers whole new possibilities for separation of rare cells which has clinically relevant prognostic value for personalized medicine.

Magnetic levitation has been an attractive tool for researchers from a variety of disciplines in life sciences ^11^ due to its simplicity, cost-effectiveness and minimal effort for processing of the sample but above all, major advantages is to provide label-free separation of target cells in a heterogeneous solution. So far, a wide range of applications including label-free separation of malaria-infected red blood cells, sickle cells ^12^, blood ^12, 13^, tumor cells ^8, 13^, adipocytes ^14^, and cardiomyocytes ^15^, realization of microparticle-based immunoassays ^16, 17^ and biofabrication of cells ^18–20^ have been achieved by applying magnetic levitation technology.

Traditionally, magnetic levitation platforms rely on bulky and costly microscope instruments in order to visualize the sampling chamber for evaluating the positions of cells ^8^. Therefore, a microcapillary glass channel containing heterogeneous cell solution is placed between two magnets with the same poles facing each other in vertical orientation and two 45° angle tilted mirrors are used on the sides of the microcapillary channel to observe the cells under a microscope. Hence, benchtop and high-cost optical microscopes are required to be operated by trained personnel in a traditional magnetic levitation configuration, which can limit its usage as a standalone device. Since tilted mirrors are used to visualize the capillary channel, effective working distance of the objectives as well as the distance between the microscope stand to objective may greatly vary. Hence, the magnetic levitation platform may not be easily compatible with various brands and models of bench top microscopes and may require additional modifications to utilize for imaging. Several studies aim to deliver portable and low-cost magnetic levitation configuration that can be adapted to a mobile phone camera for visualization. However, these studies can barely resolve features with a size of roughly a white blood cell (~20 μm) without proper illumination and additional optical configuration providing an effective magnification scheme ^21^. Thus, smartphone cameras alone may not be a fully end-to-end solution for magnetic levitation in the fields of cellular imaging or biomolecule. Alternatively, a handheld magnetic levitation platform for point-of-care testing has been shown to provide a low-cost and compact design solution that has the capability to deliver rapid analysis and ease of use, however, this design required manual focusing on microparticles that is incompatible with automated processing ^22^. As lensless digital inline holographic microscopy (LDIHM) does not utilize any optics such as lens or mirrors, LDIHM is an attractive image modality for cellular imaging and diagnostic applications and thus, the increasing number of applications has been demonstrated that LDIHM is a highly suitable choice as a low-cost and handheld cell monitoring device ^23^. The light coming from a low cost, incoherent light source (e.g. light emitting diode - LED) is spatially filtered by using a relatively large pinhole (~100 μm in diameter) to illuminate the sample that is directly placed on top of the imaging sensor with a close proximity for recording the superimposed wavefront composed of the reference wave, which is undistorted illumination light, and an object wave, which diffracts from the sample^23^. Those two waves sum up at the sensor plane to form the interference pattern also known as a hologram. Recorded hologram is required to be reconstructed to obtain the real image of the sample ^23^. Therefore, all these processes are fully digital and automated to eliminate human error and intervention for repeatable and precise measurements.

In this manuscript, we demonstrate the first use of the hybrid cell analysis platform (HologLev) composed from magnetic levitation and lensless digital inline holographic microscope. Cell densities are automatically calculated on the platform from acquired holograms of levitated cells. Holograms are processed by using the Angular Spectrum Method for reconstruction and auto-focusing algorithm, rotation, digital enhancement (e.g., adjusting brightness/contrast and binarization for mask creation) and eventually measurement of levitation heights of cells to calculate density individually automatically in a sequential order. Using HologLev, we tested viability of three different cell lines (i.e., D1 bone marrow stem cells, MDA-MB-231 breast cancer cells and U937 monocytes), differentiation of a multipotent cell line (i.e., 7F2, mouse osteoblasts) and drug response of MDA-MB-231 cells to a commonly used chemotherapy drug, Doxorubicin, by monitoring cell densities. This low-cost standalone platform requires minimal cell preparation protocols and technical expertise which would help to increase dissemination and utilization of magnetic levitation on various cell analysis scenarios.

## MATERIALS AND METHODS

### Experimental Setup

The hybrid platform was designed using computer aided design software and the main framework of the platform was manufactured by a stereolithography-based 3D printer (Formlabs Form 2), which allows precise manufacturing ^24^. After assembling 3D printed parts together, the factory built-in lens of complementary metal oxide semiconductor (CMOS) imaging sensor (Sony IMX219 - Raspberry Pi camera v2) was detached and the sensor was placed beneath of two opposing magnets (N52 grade neodymium magnets, 50 mm length × 2 mm width × 5 mm width, Supermagnete) with a separation gap less than 1 mm. Illumination and recording scheme was designed and built as a connected one piece so that the part can be slidable along the microcapillary channel for visualization of different portions of the channel. The incoming light from a light emitting diode (LED528EHP, Thor Labs, Newton, NJ) with a central wavelength of 525 nm is spatially filtered by a pinhole with a diameter of 100 μm which was manufactured by laser cutting machinery to drill the pinhole on a thin slice of aluminum sheet. Then, the light leaves the pinhole and reaches the microcapillary at the distance of about 5 cm so that plane wave approximation was ensured for further imaging processing steps ^25^.

### Magnetic Levitation Principle

Non-ionic paramagnetic solution (Gadavist®, Bayer) is utilized to levitate cell samples in the microcapillary placed between two opposing magnets in the magnetic levitation platform that eliminates the need for the use of external labelling process that provide label-free separation of cell sample with respect to their densities. This paramagnetic solution has been shown that cell viability and physiology are not adversely affected in elevated concentrations (>100 mM) ^18^.

In the magnetic levitation platform, cells tend to move to the middle point of the two magnets, where the magnetic field becomes minimum. Then, the cells become stationary, when magnetic and buoyancy forces balance each other as in the following equation^8^:

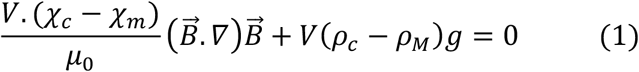

where *χ*_*c*_ and *χ*_*m*_ are the magnetic susceptibilities of cells and paramagnetic medium, *μ_0_*, is permeability of free space (vacuum) (1.2566×10^−6^ kg·m·A^−2^·s^−2^), 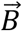 indicates the magnetic induction (*T*), *ρ_c_*, *ρ_M_* are densities of cells and medium, g is the gravitational acceleration (9.8 ms^−2^) and *V* is the volume of the cell. As seen in the equation, *V* term is cancelled out so that the levitation height profile does not depend on the cell volume. However, cells with bigger diameters come to the equilibrium levitation positions much faster. Moreover, cells have negligible magnetic susceptibility (χ_*c*_) compared to the paramagnetic solution. As a result, if we levitate the cells in the same environment, the main factor that results in unique levitation height profiles is the density of cells. In the platform, denser cells levitate closer to the bottom magnet. Hence, the cell density can be identified by measuring levitation height (i.e., the distance to the bottom magnet) of the cell.

### Image Acquisition

The entire acquisition process was performed with a Raspberry Pi 3 Model B+ by using a fully automated custom-built Python script that acquisition time and imaging parameters can be adjusted for remote and prescheduled screening and recording scheme. Once the interference pattern between the light diffracted from cells within the microcapillary channel and the undistorted background illumination was acquired, the resultant hologram was reconstructed to obtain real cell images. Thus, recorded hologram (*Ψ*_*Po*_(*x*, *y*)) on the sensor plane was back-propagated to the object plane along z-axis by using a spatial transfer function (*H*(*k_x_*, *k*_y_; *z*)) with incremental steps (*z*) to find the focused and high contrast object images by applying an embedded autofocus algorithm ^26^. Therefore, Angular Spectrum Method was used as the back-propagation algorithm and in which at each step of z, Fourier transformation of hologram was performed, and the resultant image was multiplied with transfer function ^23^. After applying inverse operation for Fourier transform, the complex-valued image was converted into amplitude and phase images (*Ψ*_*p*_(*x*, *y*; *z*))) by taken of the absolute and the argument of the complex-valued image, respectively ^23^:

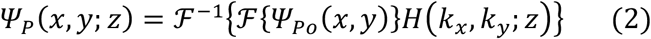

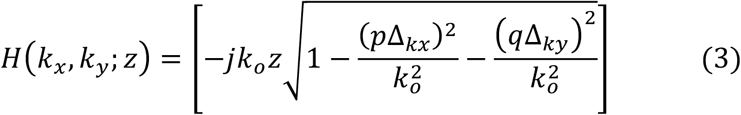

*k_o_* = *w*_*o*_/*v* is the wave number, where *w*_*o*_ is angular frequency(rad/s), *v* is the speed of the wavefront. (*x*, *y*) and (*p*, *q*) are indices for spatial and Fourier domain of the sample, respectively. Δ_*kx*_= 2π/*M*Δ_*x*_ and Δ_*ky*_ = 2π/*N*Δ_*y*_ are frequency resolutions in x and y directions (radian per unit of length), where Δ_*x*_ and Δ_*y*_ are sampling periods, and M, N are the number of samples (i.e., pixel counts), respectively.

The amplitude image was used to evaluate if the in-focus image was obtained that sharpness value of the recent amplitude image was calculated and compared against the previous image so that if the value decreased the recent image should be more in-focus than the previous one, and thus, the value and the image are updated. This process was performed within a certain interval in z direction to find optimal distance that matches with the actual object plane that raised in focus and sharp reconstructed images.

### Automated Analysis of Cell Densities from Images

Once in-focus amplitude image was obtained, Hough transformation ^27^ was applied to the image to align the bottom magnet horizontally for further density calculations in latter steps. As reconstructed holographic (i.e., amplitude) images suffer from an optical artifact presented in LDIHM accompanying with a severe irregular background noise, thresholding may become a very challenging task. Thus, a series of morphological operations were performed prior to determine the spatial coordinates of each microparticles. First, the contrast of the image was enhanced in order to relieve the edges which were suppressed by the noise coming from overlapped twin image. For a binary mask, the threshold was tuned to identify cells with certain diameters. After creation of a binary mask that indicates the positions of each cell, the center of gravity of each blob (i.e., cell mask) was detected. Accordingly, the vertical distances between calculated coordinates of each blob and the horizontal plane indicating the upper edge of the bottom magnet whose height was automatically determined after Hough transform were determined ^28^. Note that calculated horizontal lines using Hough transform were also used to determine the microcapillary channels’ top and bottom contours. Hence, any particle found higher or lower than this region was excluded before density calculations. Therefore, levitation heights of cells were converted into density values by using the calibration curve of polystyrene microspheres with known densities. All computational processes were performed by custom built script using MATLAB 2018a.

### Levitation of Polyethylene Microspheres

Polyethylene microspheres with different densities, 1.00 g mL^−1^ (with size of 10-20 μm), 1.02 g mL^−1^ (with size of 10-20 μm), 1.05 g mL^−1^ (with size of 45-53 μm) and 1.09 g mL^−1^ (with size of 20-27 μm) (Cospheric LLC., ABD), were levitated in Phosphate Buffered Saline (PBS, Gibco) with 1% (w/v) Pluronic F-127 (Sigma Aldrich, Germany) containing 25 mM, 50 mM, 75 mM, and 100 mM Gadolinium (Gd^3+^) (Gadavist®, Bayer). Levitated heights of microspheres were measured in the platform after they reached stationary position in 10 minutes.

### Cell Culture

D1 ORL UVA (mouse bone marrow stem cells) and MDA-MB-231 (breast cancer cells) were cultured in Dulbecco’s modified Eagle’s high glucose medium (DMEM, Gibco) supplemented with 10% fetal bovine serum (ECS0180, Euroclone) and 1% penicillin/streptomycin (Penicillin/Streptomicin 100X, Euroclone). U-937, human monocyte cells were cultured in a Roswell Park Memorial Institute medium (RPMI-1640, ECM0495L, Euroclone) containing 10% FBS and 1% penicillin/streptomycin. The cells were grown in a humidified 37 °C incubator with 5% CO_2_. The growth medium was changed every other day and the cells were passaged every four to six days.

7F2 (mouse osteoblasts) cells were grown in a growth medium supplemented with %10 FBS and 1% penicillin/streptomycin at 5% CO2 humidified atmosphere at 37 °C. 7F2 cells were cultured in alpha modified essential medium (αMEM, Sigma Aldrich). The growth medium was refreshed every 2–3 days and the cells were passaged every 4–6 days. For differentiation of the 7F2 cells into adipocytes, the cells were induced in αMEM with induction agents for 10 days ^14^. The adipogenic induction medium was refreshed every 2–3 days.

### Cell Preparation for Density Analysis in HologLev

Live D1 ORL UVA and MDA-MB-231 cells were trypsinized, spun down at 125× g for 5 minutes and the supernatant was discarded. U-937 cells were prepared by using the same protocol without trypsinization. The cells were resuspended to 10^5^ cells per mL in the culture medium with 25 mM or 100 mM Gd^3+^, then 40 μL of cell solution was loaded into the capillary channel. After 10 minutes of levitation in HologLev, they were imaged with integrated LDIHM system.

The levitation profile of dead D1 ORL UVA, MDA-MB-231, and U-937 and were also investigated at 25 and 100 mM Gd^3+^. To this end, dead cells are obtained by introducing 1:1 (v/v) dimethyl-sulfoxide (DMSO, Sigma Aldrich) to cell mediums. Then, cells were resuspended to 10^5^ cells/mL concentration in PBS containing (w/v) 1% Pluronic F-127 with 25 mM or 100 mM Gd^3+^. Then, 40 μL of cell sample was loaded into the capillary in the platform and inspected after 10 minutes of levitation.

Mouse bone marrow osteoblast 7F2 cells that were cultured in the growth and adipogenic induction media were trypsinized at the 15^th^ day and were centrifuged at 125× g for 5 minutes. Then, culture medium was removed, and the pellet was resuspended to 10^5^ cells per mL in the culture medium containing 25 mM Gd^3+^. 40 μL sample was loaded into the microcapillary channel for inspection after 10 minutes of levitation.

### Doxorubicin Drug Response Experiments

MDA-MB-231 cells were trypsinized and consecutively spun down at 125× g for 5 min to discard supernatant. The cells were resuspended to 10^5^ cells per mL in the culture medium. Afterwards, Doxorubicin chemotherapy drug, which has IC50 concentration of < 6 μM ^29, 30^ for MDA-MB-231 cells, were directly added to the growth medium at 4.2 μM, 8.4 μM and 16.8 μM concentrations. Then, 40 μL of cell solution with 100 mM Gd^3+^ was loaded into the capillary channel. The cells were levitated in HologLev, which was placed within a cell incubator whose temperature was set to 37 °C. Cells were monitored in the platform to assess the effect of drug treatment for 24 hours.

### Statistical Analysis

Live and dead cells of D1 ORL UVA, U-937, and MDA-MB-231 were analyzed using an unpaired two-tailed t-test. For cell differentiation analysis, both cell densities and skewness of densities were compared with an unpaired t-test. The lowest 5^th^ percentile of densities of differentiated and undifferentiated cells was also compared with a two-tailed t-test. *, *** and **** represent p<0.05, p<0.001 and p<0.0001, respectively.

## RESULTS AND DISCUSSION

### Experimental Setup

In this work, the use of a miniaturized lensless microscopy-integrated magnetic levitation platform, HologLev, to monitor and analyze different cell lines was demonstrated by combining two separate technologies to provide low-cost, easy-to-use, and portable cell separation analysis (Figure 1a and Figure S1). Hybrid platform was enclosed in a 3D printed framework that was mainly composed of illumination and acquisition scheme as a slidable one piece and magnetic levitation section that involves two opposing magnets and microcapillary channel in between (Figure 1a). Once the sample was placed in the microcapillary between magnets in the platform, heterogeneous cell solution was levitated, and cells reach their equilibrium position when magnetic and buoyancy forces acting on each cell are in equilibrium (Figure 1b). Full field of view (FOV) acquired from the proposed platform serves far more wider sampling area compared to that of fluorescence image captured with 10× microscope objective on an inverted fluorescence microscope for viability assay using Calcein AM and Propidium Iodide on MDA-MB-231 cells (Figure 1c). This also illustrated the compatibility of our platform to simply be integrated to a fluorescent microscope by placing a mirror in front of the LED/pinhole scheme for fluorescent acquisition of the same portion of the microcapillary followed by hologram acquisition (Figure S2).

**Figure 1.**
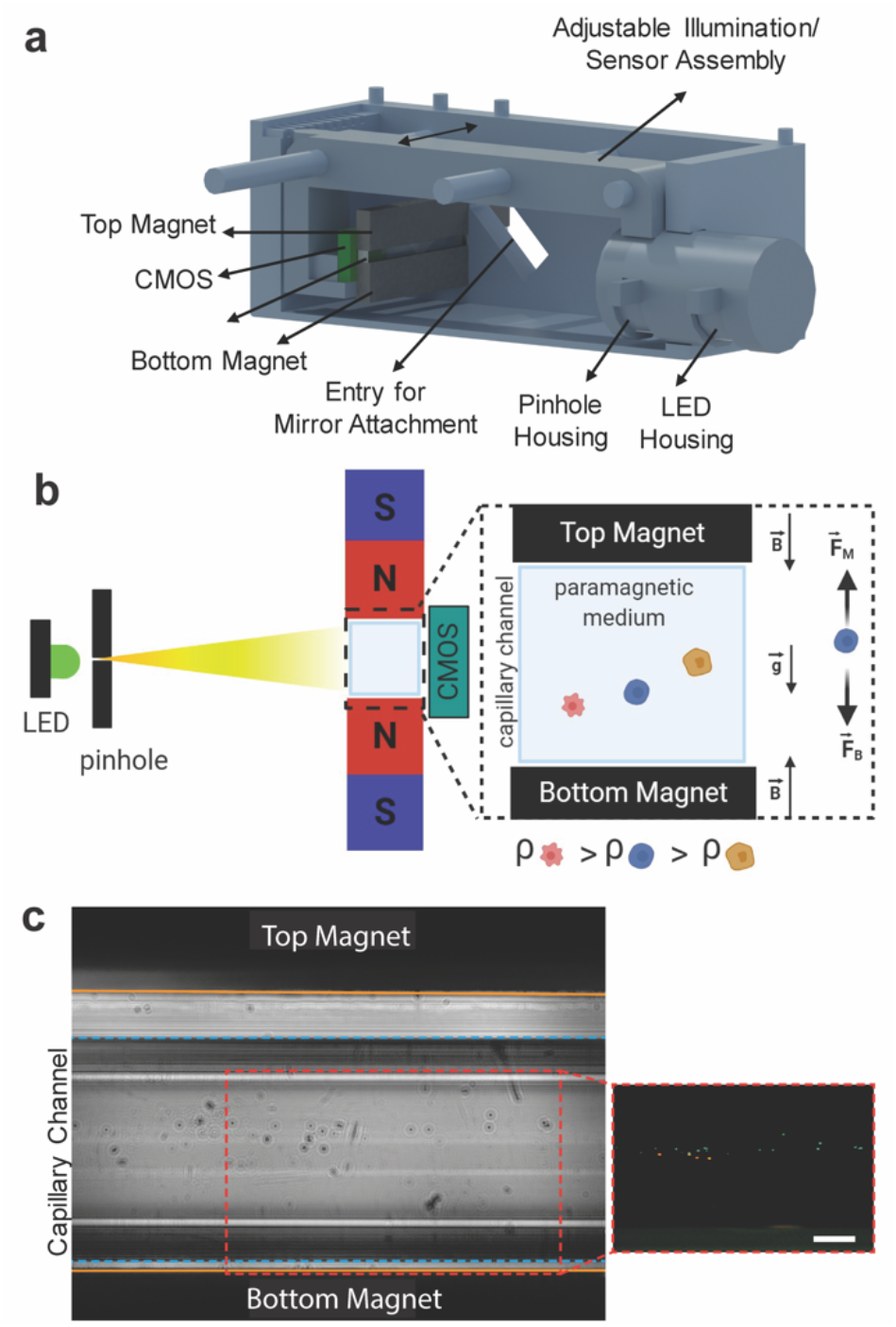
Principles of HologLev. (a) Illustration of the cross-sectional view revealing interior components. (b) Illustration of magnetic levitation principle. Particles (or cells) dissolved in a paramagnetic liquid inside the platform come to a unique equilibrium point and levitate under the balance of magnetic force 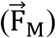 and buoyancy force 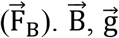, and ρ represent magnetic induction, gravity, and density, respectively. (c) Comparison of the field of the view between the holographic microscope and fluorescence microscope images of live (green) and dead (red) MDA-MB-231 cells using the mirror attachement in the platform showing magnets (orange lines) and microcapillary channel (blue dashed lines). Scale bar: 200 μm.

The microspheres and cells residing in the microcapillary channel were analyzed in terms of their levitation heights (i.e., their distance from the top surface of the bottom magnet) under the external magnetic field. After diffraction of the light from the microspheres/cells within the microcapillary channel, the interference pattern composing of the undistorted reference wave and the diffracted object wave was recorded by CMOS imaging sensor at a distance less than a millimeter. Since there were no optics used between the sample and the imaging sensor, the diffracted light from the sample was placed as close as possible to the imaging sensor to acquire hologram without the loss of image quality. The spatial resolution was also limited with the pixel size of the imaging sensor (i.e., 1.12 μm). We measured the spatial resolution of our system by imaging a standard optical resolution chart USAF 1951 (Figure S3). Our hybrid platform was capable of resolving features that match with group: 8, element:1 of USAF 1951 corresponding to 1.95 μm which was adequate for imaging of mammalian cells whose diameters are greater than the resolved feature size. Thus, quantitative assessment of spatial resolution indicates that the platform cannot only have portable and low-cost design but is also capable of delivering high resolution images that can compete with high cost and bulky benchtop microscopes for a wide range of applications from cell imaging to point-of-care-testing.

Once cells became levitated in the platform, the hologram of the sample was acquired (Figure 2a). The hologram was reconstructed by Angular Spectrum Method and auto-focusing algorithm (Figure 2b, Video 1). Afterwards, the reconstructed image was enhanced digitally and rotated for aligning the bottom magnet horizontally (Figure 2c). After successfully completing of these steps, vertical distances (blue line) between bottom magnet (green line) and each cell coordinates of center of gravity were calculated for determining relative levitation heights that were then converted into density using the calibration curve (Figure 2d).

**Figure 2.**
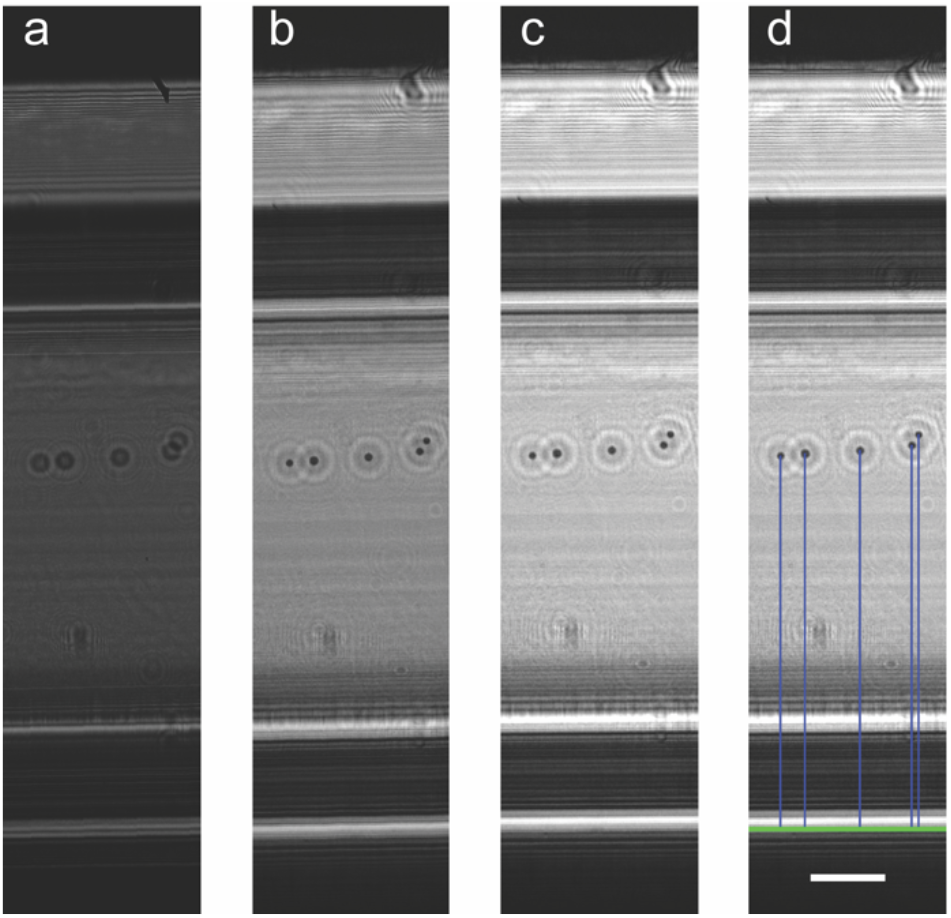
Illustrations of digital image processing steps through determination of relative levitation heights of MDA-MB-231 cells spiked in 100 mM Gd^3+^. (a) Recorded cell hologram, (b) back-propagated amplitude image, (c) enhancement and rotating of reconstructed image, (d) detected cells regarding their relative levitation heights (blue lines) with respect to bottom magnet (green line). Scale bar: 200 μm.

### Calibration of the platform

For cell experiments in HologLev, calibration of the platform was done using polyethylene standard density microspheres (i.e., 1.005, 1.026, 1.050, and 1.090 g/mL). Due to their diamagnetic properties, they migrate from the denser magnetic field regions (i.e., from top and bottom magnets) and levitate at unique heights depending on their density under the balance of magnetic and buoyancy forces. The levitation heights of microspheres were determined automatically from the reconstructed images. Calibration curves obtained with the microspheres’ images under different Gd^3+^ concentrations were presented in Figure 3. Density versus levitation height was determined using the standard equations created by fitting the experimental levitation data into linear curves. According to the results, increased Gd^3+^ concentrations caused microspheres levitate at higher heights and the slope of calibration curves was decreased and so the sensitivity of density analysis. The results were also analyzed by manual counting using Cell Counter Plugin in Image J (Fiji). According to the results, the calibration curve obtained by manual analysis is in good agreement with the calibration curve obtained using automated digital image analysis (Figure S4).

**Figure 3.**
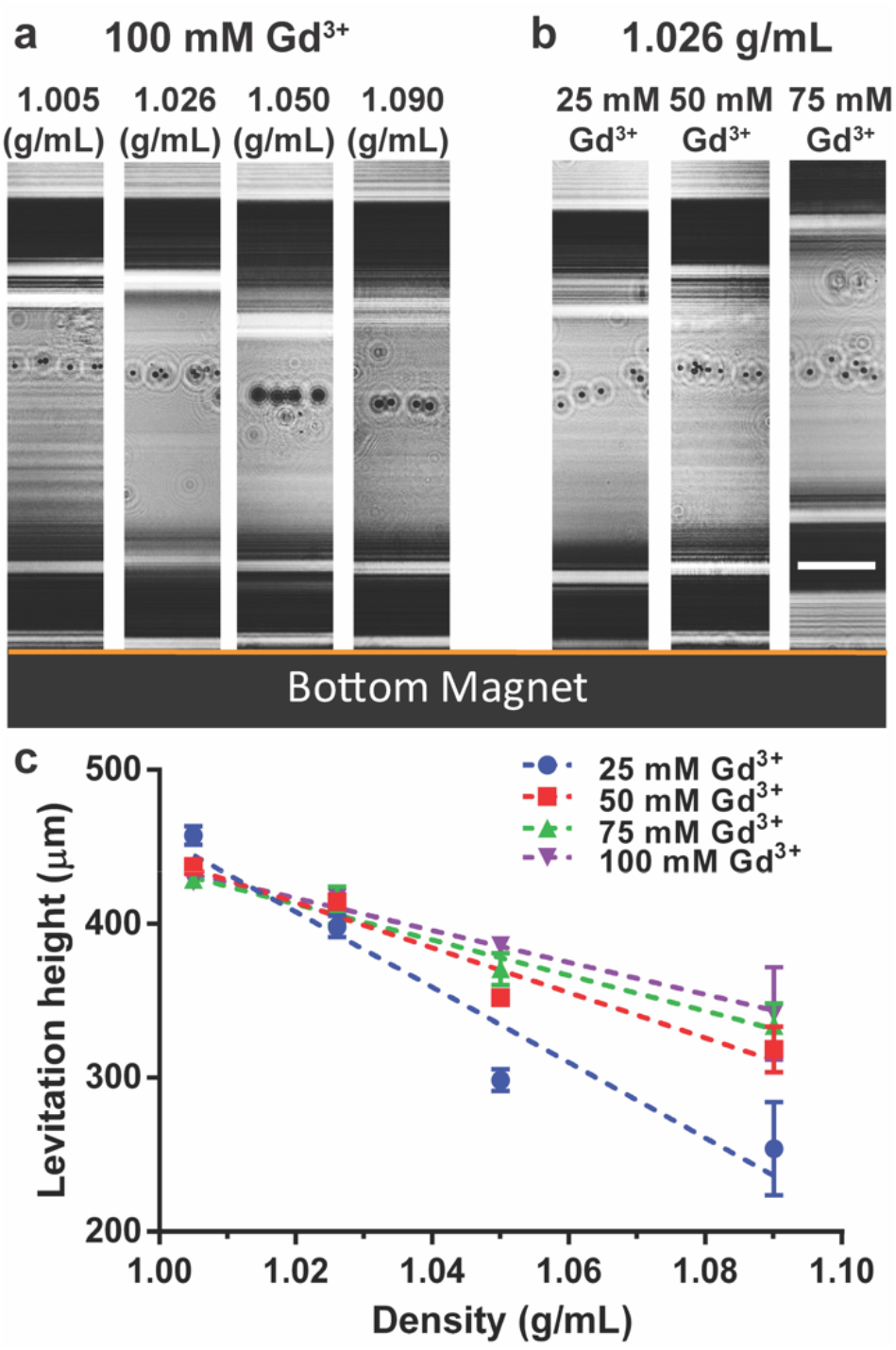
Calibration of HologLev using density-standard microspheres at 1.005, 1.026, 1.05 and 1.09 g/mL densities. (a) Reconstructed images of standard density microspheres at 100 mM Gd^3+^ concentrations. (b) Reconstructed images of 1.026 g/mL microspheres at 25 mM, 50 mM, and 75 mM Gd^+3^ concentration. Scale bar: 200 μm. (c) Levitation height analyses of different density microspheres under tested Gd^3+^ concentrations (i.e., 25, 50, and 75 mM). Dashed lines represent linear fitting of density versus levitation height data. R^2^ values for 25, 50, 75, and 100 mM Gd^3+^ experiments are 0.93, 0.95, 0.98, and 0.99, respectively. The data represent as mean ± standard deviation of three different experiments.

### Magnetic Levitation of D1 ORL UVA, U-937, and MDA-MB-231 Cells

Live and dead cells of three different cell lineages (D1 ORL UVA, U-937, and MDA-MB-231) were analyzed within microcapillary using magnetic levitation principle. Dead cells of those cell lines were located at the bottom of the capillary channel when a 25 mM Gd^3+^ was used in the levitation medium. To be able to levitate dead cells at higher heights, 100 mM Gd^3+^ was tested (Figure 4a). We measured densities of D1 ORL UVA, U937 and MDA-MB-231 cells at 100 mM Gd^3+^ for live as 1.0721 ± 0.0065, 1.0680 ± 0.0064 and 1.0660 ± 0.0110 g/mL, and for dead as 1.1653 ± 0.0087, 1.1723 ± 0.0054 and 1.1681 ± 0.0076 g/mL, respectively. At 100 mM Gd^3+^, the densities of live and dead cells of the same cell line were statistically different when a t-test was applied for each of three cell lines (Figure 4b). In this section, cell death was provided with the introduction of an amphipathic solvent, DMSO, which has cytotoxic effects on cells ^31^. DMSO increases cell permeability by interacting with the cells’ plasma membrane ^32^. As cell death occurred due to DMSO, disintegration of cellular structure probably caused accumulation of Gd-based medium inside the cell. Hence, the effect of cytoplasm on cell density was eliminated, and the remaining cell debris had a higher density than the intact cell. To this end, dead cells were collected at lower heights than the live ones. These results showed that change in cell density due to the loss of viability can be effectively identified using HologLev, based on monitoring densities of cells in the channel. For instance, cell death is essential for maintaining homeostasis, however, any disorder of this mechanism can indicate severe diseases. Vascular pathology resulted in significant death in endothelial and vascular smooth muscle cells can exhibit a vital vessel remodeling that can indicate such diseases like atherosclerosis and formation of aneurysm^33^. Thus, rapid, low-cost, and easy identification of death cell populations in blood plasma using this hybrid device can offer early diagnosis and preventive treatments for variety of cardiovascular abnormalities in patients who suffer from a severe condition which can be unlikely to be determined without any invasive (e.g., angiography) or advanced radiological imaging techniques (e.g., computer assisted tomography) in present clinical routine.

**Figure 4.**
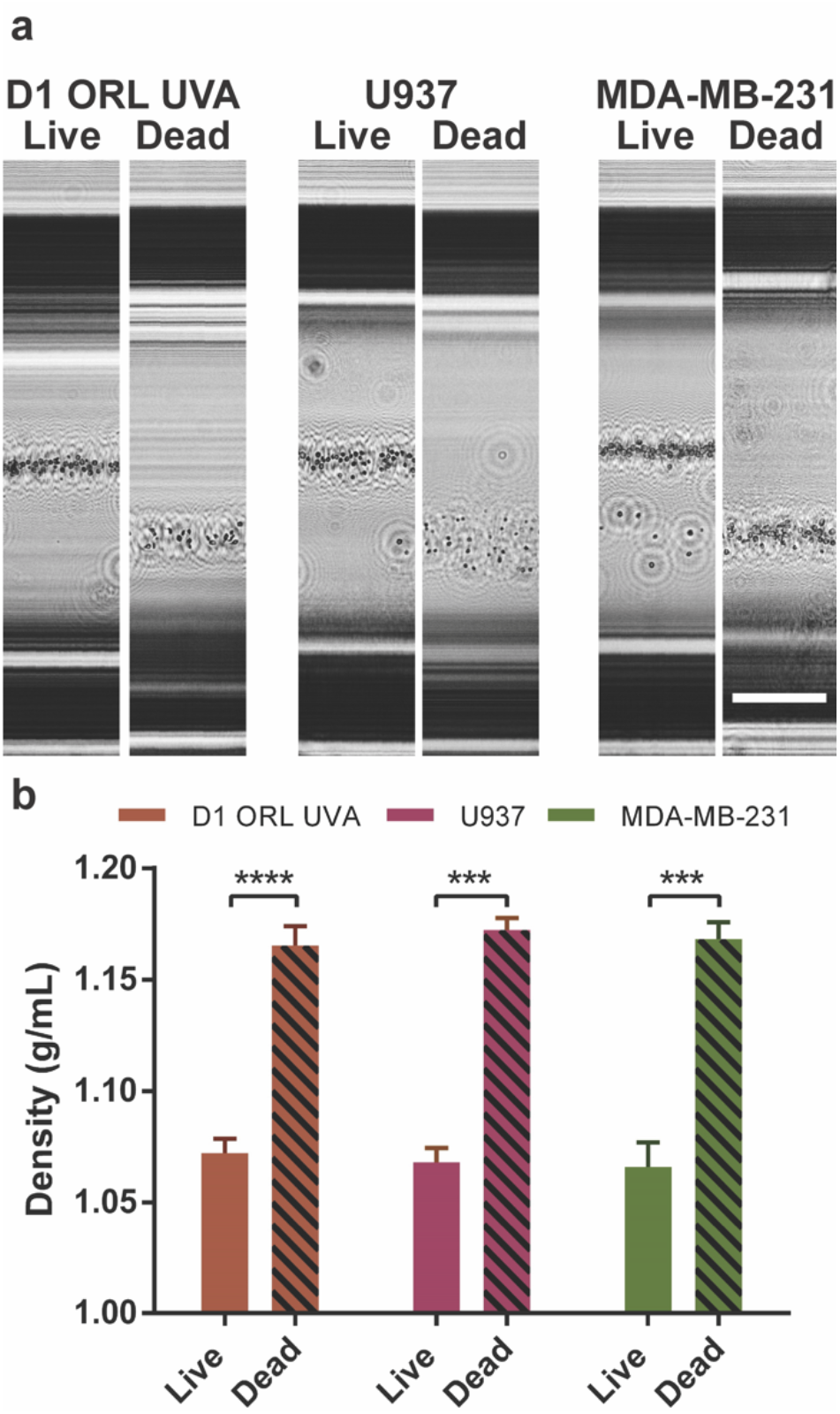
Density analysis of live and dead cells. (a) Reconstructed images and (b) calculated average densities of live and dead D1 ORL UVA, U-937 and MDA-MB-231 cells spiked in 100 mM Gd^3+^. The data represent as mean ± standard deviation of 3 different experiments. The data are compared using an unpaired t-test. *** and **** indicate p<0.001 and p<0.0001, respectively. Scale bar: 200 μm.

### Monitoring Adipogenic Cell Differentiation

Assessment of stem cell differentiation is of great interest in the field of regenerative medicine due to therapeutic potential of stem cells for a variety of diseases ranging from spinal cord injuries to cancer^34^. Thus, it could provide significant advantages to deliver a rapid, low-cost, and portable platform to evaluate differentiated cells from the rest of the population with minimal need for sample processing and expertise. In a previous study, adipogenic differentiation of bone marrow origin has been reported using microscope-based magnetic levitation system ^14^. To demonstrate our platform’s capability to distinguish cell differentiation, similarly we used 7F2 cells which were treated to adipogenic differentiation using adipogenic induction medium whereas undifferentiated 7F2 cells were maintained in growth media as control. 7F2 cells were allowed to differentiate by accumulating lipid content within their cytoplasm which essentially affects the change in density between undifferentiated and differentiated cell populations subject to treatment. Furthermore, adipogenic differentiation of cells exhibit distinct morphological tracks accompanying density change. After the 15^th^ day of cell growth in the culture media for differentiation of 7F2 cells, differentiated and undifferentiated (control) cell solutions were prepared and Gd^3+^ solution corresponding to 25 mM was added to both cell solutions prior to detection of differentiation experiments with our platform. Within 10 minutes after loading microcapillary channels containing solutions, cells reached equilibrium positions and holographic images of each cell group were acquired. The positions of cells in the microcapillary channel revealed the average densities of differentiated cells resided similar to undifferentiated (control) cells (Figure 5a and 5b). However, it seems like differentiated cells occupy a far broad range of levitation heights indicating a more distributed density value (Figure 5c) and undifferentiated cells were levitated a tighter band at the upper portion (5^th^ percentile) of the positions of differentiated ones (Figure 5d). Thus, it can be deduced that differentiation induces decrease in density and this change dictates the cell to shift towards higher levitation heights. In conclusion, we demonstrated that cell differentiation can be successfully detected with a simple, low-cost, and portable setup that does not require any trained personnel or sophisticated and expensive/time consuming fluorescence labelling protocol from only acquisition and analyzing of holographic images.

**Figure 5.**
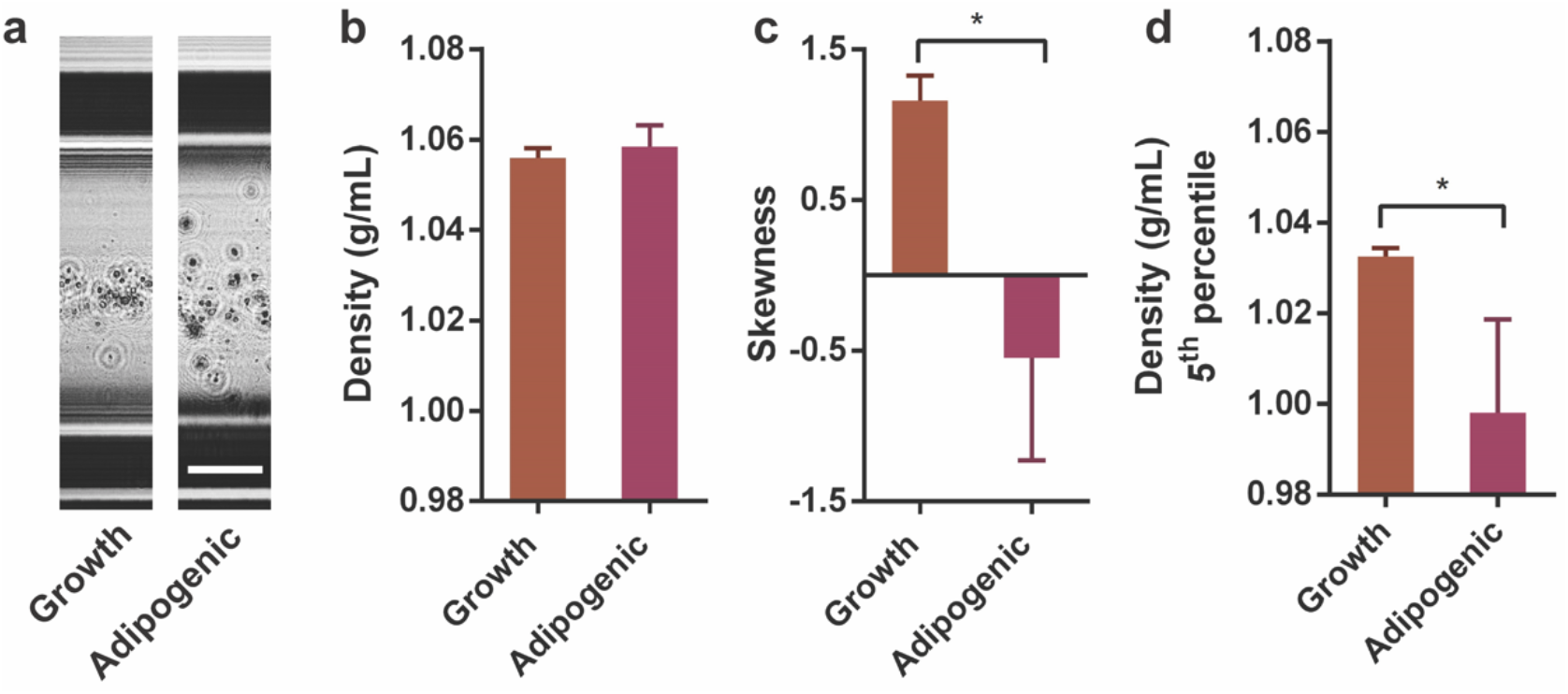
Determination of adipogenic cell differentiation. (a) Reconstructed images of 7F2 (mouse osteoblasts) that were cultured in the growth (7F2G) and adipogenic (7F2A) induction media levitated in 25 mM Gd^3+^ solution. Comparison of (b) average densities, (c) skewness of density for distribution analysis, (d) the lowest 5^th^ percentile of densities. The data represent as mean ± standard deviation of three different experiments. The data are compared using an unpaired t-test. * indicates p<0.05. Scale bar: 200 μm.

### Monitoring of Drug Response of Cancer Cells

Measuring drug response is one of the most important and widely studied fields mainly for drug delivery and cancer treatment. So, an effective drug is expected to deliver appropriate host response without any adverse side-effects while delivering active molecules to the target biological system. However, it is challenging to measure these effects *in vivo,* thus, a variety of cell assays are used to assess drug effects on cells by measuring viability, toxicity etc. Hence, these assays should be deployed with ease and low-cost. To provide our systems capability for real time drug response monitoring, we conducted experiments with MDA-MB-231 breast cancer cells treated with a common chemotherapy drug, Doxorubicin. We tested three different concentrations of the drug (i.e., 4.2 μM, 8.4 μM, and 16.8 μM) on cancer cells and analyzed change in density of cells during a period of 24 hours along with non-treated cells in a cell incubator. In the beginning of the experiments for each dose, cells were well-separated and can be distinguishable at single cell level from acquired holographic images. Although cells became aggregated in time, they remained levitated in the capillary. It is challenging to backpropagate the holographic images because the twin image artifact suppresses the fine details to reconstruct the cluster formation. Therefore, we present raw holographic images to demonstrate the change in levitation heights with increasing drug dose (Figure 6a). We observed that 24 hours drug treatment altered the levitation height so the density profile of the cells (Figure 6b). Increasing dose, which implies reducing viability of cells, boosted the average density of the cells. Cell densities after 24 hours were measured as 1.0621, 1.1608, 1.1531 and 1.2415 g/mL for control, 4.2 μM, 8.4 μM and 16.8 μM, respectively, being higher than the control. For all drug-treated cells, the 24^th^ hour densities of the cell clusters were shown higher densities of live MDA-MB-231 cells shown in Figure 4b. We additionally tested any significant variation of growth medium during 24 hours of experiment to manipulate the change in levitation heights of treated cells that may misguide the effect of the drug treatment. Therefore, we used polymeric microspheres with a known density (i.e., nominal density of the microsphere that manufacturer indicated is 1.05 g/mL) and levitated them for 24 hours in a cell incubator at 37 °C in which the temperature was set the same as in the drug treatment experiments (Figure S5). We verified that levitation height of microspheres was constant for 24 hours which implies that density change of the medium due to heat and humidity in the incubator is negligible. Consequently, our hybrid platform successfully reveals the density changes in drug treated cancer cells in cell culture environment to provide a potential benefit for use in crucial clinical applications without any labelling and expertise for testing drug efficacy while delivering fully automated imaging-based solution.

**Figure 6.**
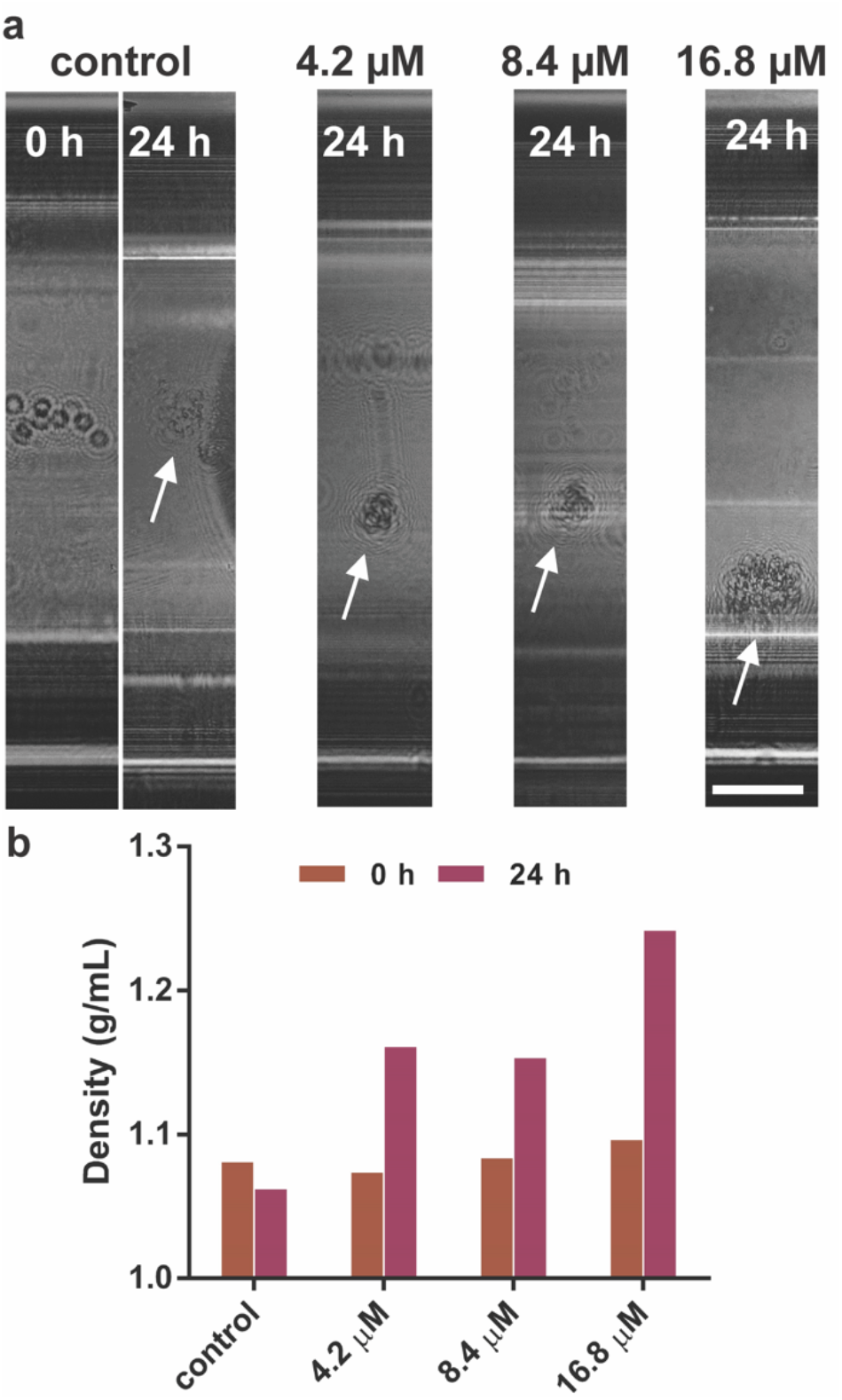
Drug response of levitated MDA-MB-231 cells in 100 mM Gd^3+^ treated by Doxorubicin with different doses of 4.2 μM, 8.4 μM and 16.8 μM. (a) Holographic images and (b) cell densities for different drug doses. Cells clusters were indicated as white arrows. Scale bar: 200 μm.

### Density-Based Analysis

Density can be considered as an important physical biomarker to differentiate different cell populations ^6, 7^. Moreover, the change in cells’ mass to volume ratio may reflect certain cellular processes like cell death ^35^, cell differentiation ^36^ and state of disease ^37^. In this work, we tested our novel platform for density-based label-free magnetic levitation applications that aim to monitor cells at the single-cell level. Unlike other common single cell analysis methods (i.e., flow cytometry) that require time-consuming and costly labeling protocols of the target cells with additional molecules such as fluorescent antibodies ^38^ or quantum dots ^39^, our platform utilized only the intrinsic label-free biophysical properties of cells. The one of the workhorse single-cell sorting technique, so-called fluorescent-activated cell sorting (FACS) offers tremendous sensitivity for most of the cells with a high throughput (>10.000 cells/s) and provides multiple parameter analysis ^40^. Yet, FACS has bottlenecks related to instrumentation costs and the amount of materials used per analysis ^41^. Moreover, cell labeling using fluorophore-conjugated antibodies may cause loss of cell viability ^42^. On the contrary, magnetic levitation-based tests in HologLev enables cell assays done in a portable, low-cost, label-free and automated manner without affecting the cell viability for the Gd^3+^ concentrations used in this work (i.e., 25-100 mM) ^18^. Reconstructed holographic images heavily suffer from an artifact called twin image so that the cell concentration should be sufficiently sparse in order to get well-recovered binary masks to calculate the spatial positions of each microparticle. Since the recent automation framework is sensitive while determining the magnet and capillary positions to exclude irrelevant particles followed by computations of levitation heights, future studies will focus on developing an adaptive and modified thresholding algorithm for holographic imaging which is independent from localized noise around microparticles. Traditional magnetic levitation-based cellular analysis platforms have required external imaging equipment (e.g., benchtop microscopes) which not only limits the platform’s capability of testing in clinical and home use but also dramatically increases the overall cost of the tests. Thus, we present, the first time, a standalone maglev platform with built-in imaging modality that provides high resolution images (<2 μm) without need for peripheral optics, auxiliary hardware and manual optical focusing ^43^. The entire platform can be built less than $100 by using widely accessible consumable and with intermediate experience in electronics and optics (Table S1). The most expensive components of the platform are imaging sensor and the microcomputer used for image acquisition which can be replaced by more compact and low-cost solutions in the oncoming studies. In addition, the platform is aimed to be a handheld and multimodal imaging microscope so that it can retrieve information from different domains and thus, HologLev is capable of performing fluorescent imaging alongside with holographic imaging by placing the platform on a stand of the fluorescent microscope as if it is a regular specimen. Thus, the next generation of HologLev could be equipped with built-in fluorescent lights source and imaging scheme. This system could be used as a cell cytometer in the future by integrating pumping functionality in this standalone platform. In addition, our platform can also be used for analysis of polymer microspheres that can be applied for point-of-care biomarker quantification by using antibody decorated microspheres^16^.

## CONCLUSION

Magnetic levitation technique based on microscopic examination for determining levitation heights of microparticles or cells depends on bench-top microscopes which are composed of highly fragile and costly optics along with dependency of trained personnel to operate that limits their usage. In this work, we demonstrated the possible use and practical advantages of the novel hybrid magnetic levitation based LDIHM platform in life sciences for a variety of applications including cell viability, stem cell differentiation and drug testing. Our novel hybrid magnetic levitation platform showed that it can deliver high resolution images with a quantitative and fully automated pipeline. The standalone platform has low-cost fabrication cost that can disseminate usage of magnetic levitation technology in different applications. It provides end-to-end automated image acquisition and processing algorithms which can be further extended to remote and real-time screening of cells to track cellular movement and morphological changes.

## SUPPORTING INFORMATION

See the supporting information for Figures S1-S5, Table S1 and Video 1.

## Acknowledgements

Authors would like to thank The Scientific and Technological Research Council of Turkey (119M052) for funding this work. K.D. and O.S. acknowledge the support of Turkish Council of Higher Education for 100/2000 CoHE doctoral scholarship. K.D is thankful for the helpful discussions with Ersin Cine from Department of Computer Engineering at IZTECH while developing the automation framework and with Ali Aslan Demir from Department of Photonics at IZTECH while improving the framework. H.C.T. would like to thank RPh. Vecdi Tekin from Mudanya Pharmacy (Bursa, Turkey) for supplying Doxorubicin.

## Author Contributions

H.C.T. and E.O. conceived and designed the study; K.D. designed and fabricated the platform, and developed the automation framework; K.D, S.Y and E.Y. performed experiments; S.Y. performed statistical analysis; M.A. and O.S. performed cell culture, K.D., S.Y., E.O. and H.C.T. analyzed the data; K.D., S.Y., E.Y., E.O. and H.C.T. wrote the manuscript.

