## supporting information for "HologLev: Hybrid Magnetic Levitation Platform Integrated with Lensless Holographic Microscopy for Density-Based Cell Analysis"

Kerem Delikoyun[1], Sena Yaman[1], Esra Yilmaz[1], Oyku Sarigil, Muge Anil-Inevi[1], Engin Ozcivici[1], H. Cumhur Tekin[1, 2*]

[1]Department of Bioengineering, Izmir Institute of Technology, Izmir 35430, Turkey

[2]METU MEMS Center, Ankara 06520, Turkey

† K.D and S.Y. contributed equally to this work and are listed alphabetically.

**Table of Contents**

Figure S1 | Illustration of HologLev including Raspberry Pi for image acquisition.....S2

Figure S2 | Illustration of multimodal imaging capability of hybrid platform.....S3

Figure S3 | Spatial resolution achieved by hybrid platform using USAF 1951 optical test chart.....S4

Figure S4 | Levitation analysis of microspheres using manual and automated counting for calibration of the platform under 25 mM $Gd^{+3}$.....S5

Figure S5 | Levitation height change of microspheres with an average density of 1.05 g/mL in an incubator platform for 24 h.....S6

Table S1 | The prices of the components used for building the platform.....S6

Video 1 | The video of hologram (left) and back-propagated amplitude image (right).......S7

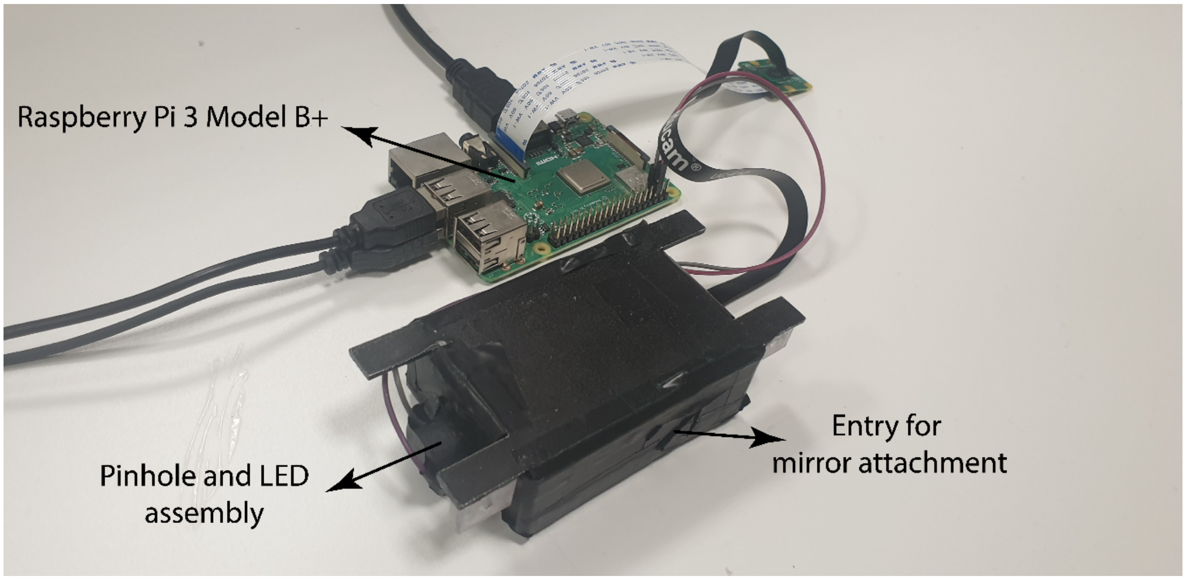


**Figure S1.** Illustration of HologLev including Raspberry Pi for image acquisition.

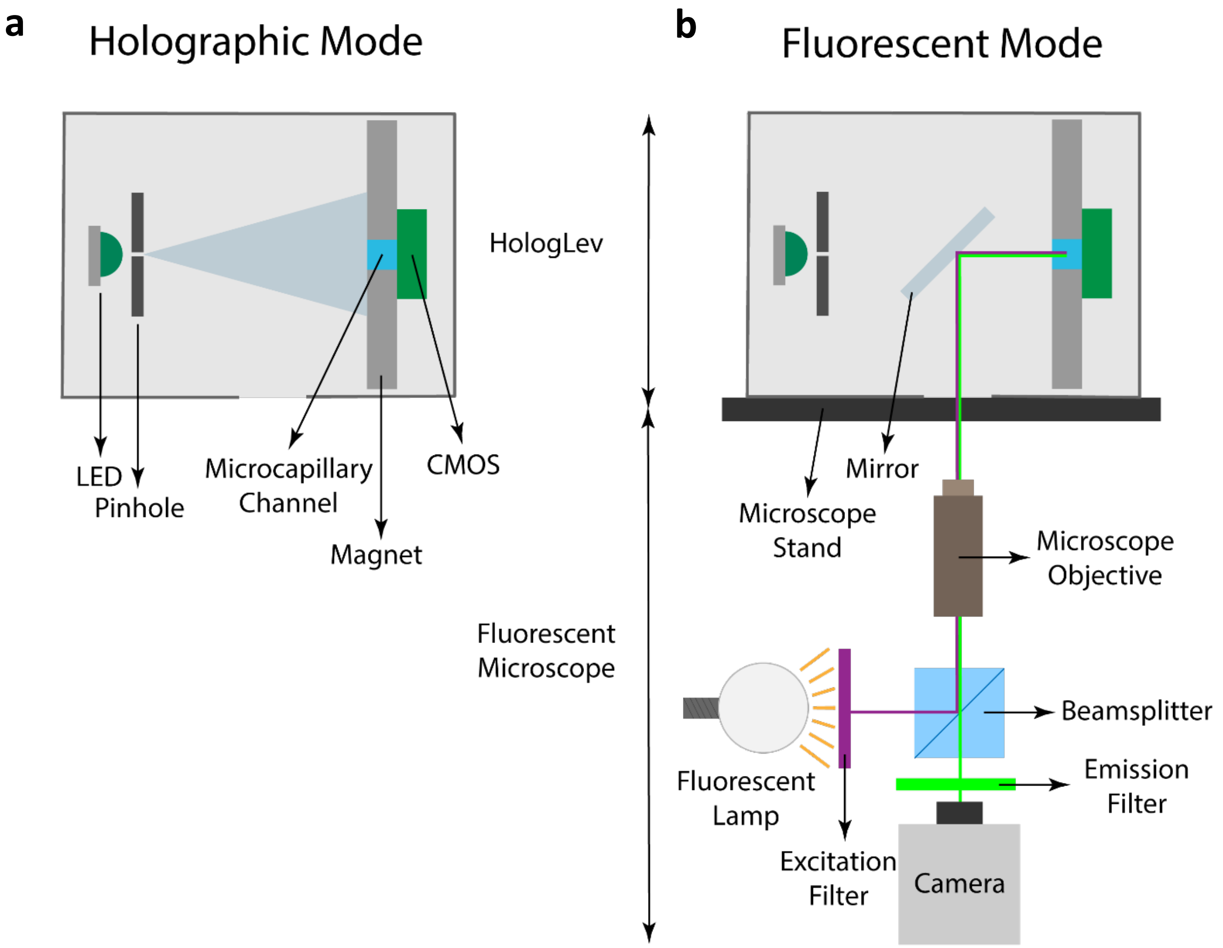


**Figure S2.** Illustration of multimodal imaging capability of hybrid platform. (a) Holographic mode operated with LDIHM scheme and (b) Fluorescent mode operated with a fluorescent microscope (Zeiss Axio Vert A1 with Zeiss Colibri 7 Type RGB-UV LED fluorescent illumination and Zeiss filter set FS 90 HE) by placing a tilted mirror between illumination scheme of LDIHM and the magnet assembly from mirror entry opening shown in Figure S1.

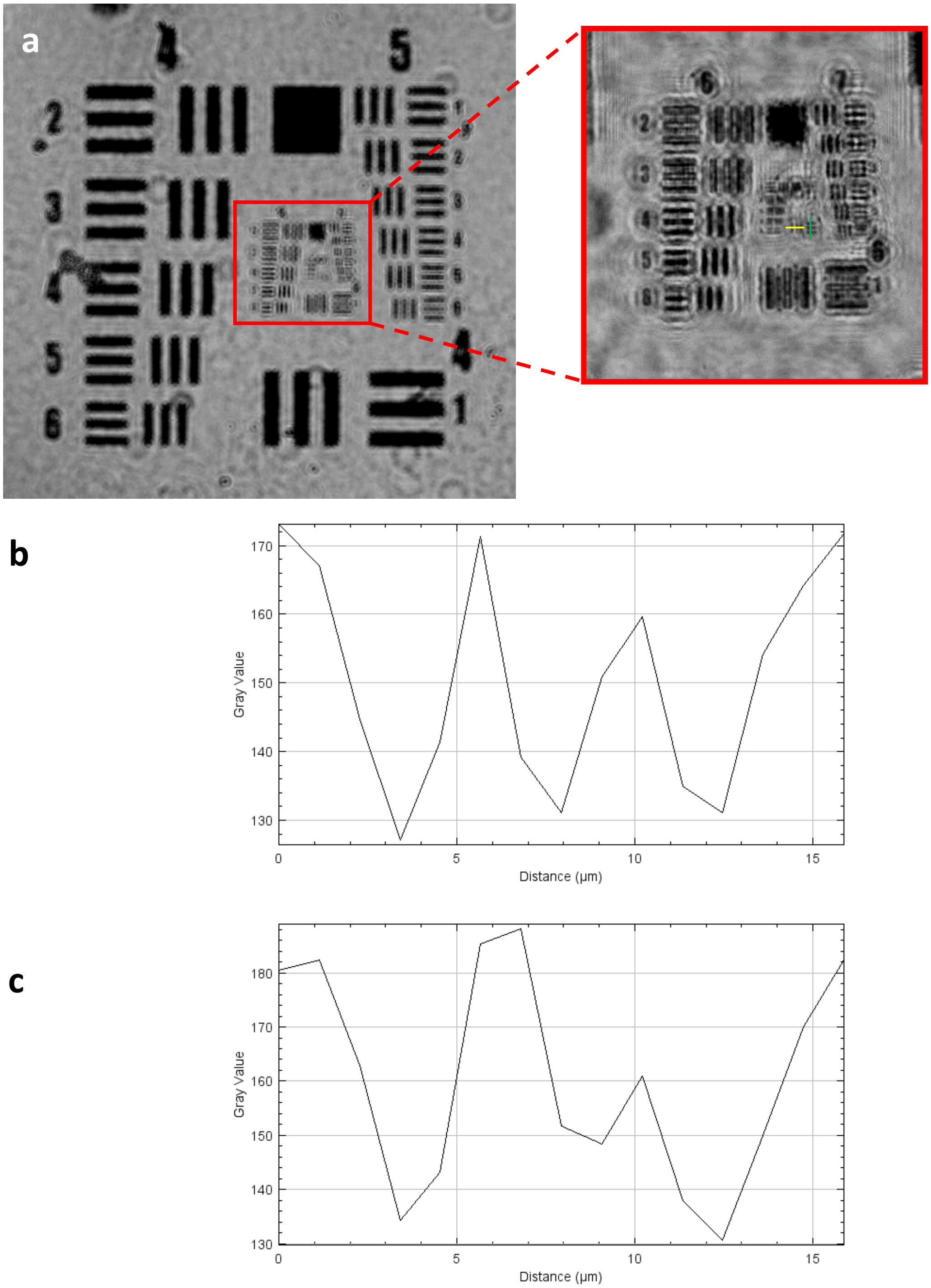


**Figure S3.** Spatial resolution achieved by hybrid platform using USAF 1951 optical test chart. (a) Reconstructed image of test chart and magnified inner portion including group 8 and 9. (b, c) Intensity profile of vertical bars (green line) and horizontal bars on group:8 – element:1 (yellow line), respectively.

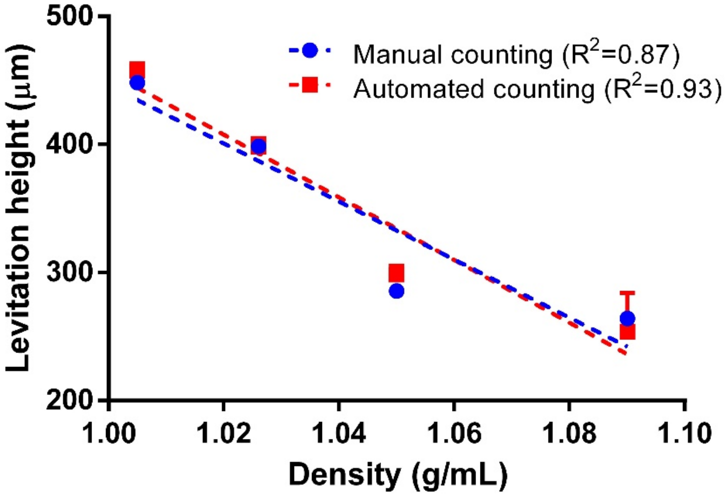


**Figure S4.** Levitation analysis of microspheres using manual and automated counting for calibration of the platform under 25 mM $Gd^{+3}$. The standard equations giving levitation height (h, μm) vs. density (d, g/mL) are h= -2266d + 2712 and h= -2448d + 2904 for manual and automated counting, respectively.

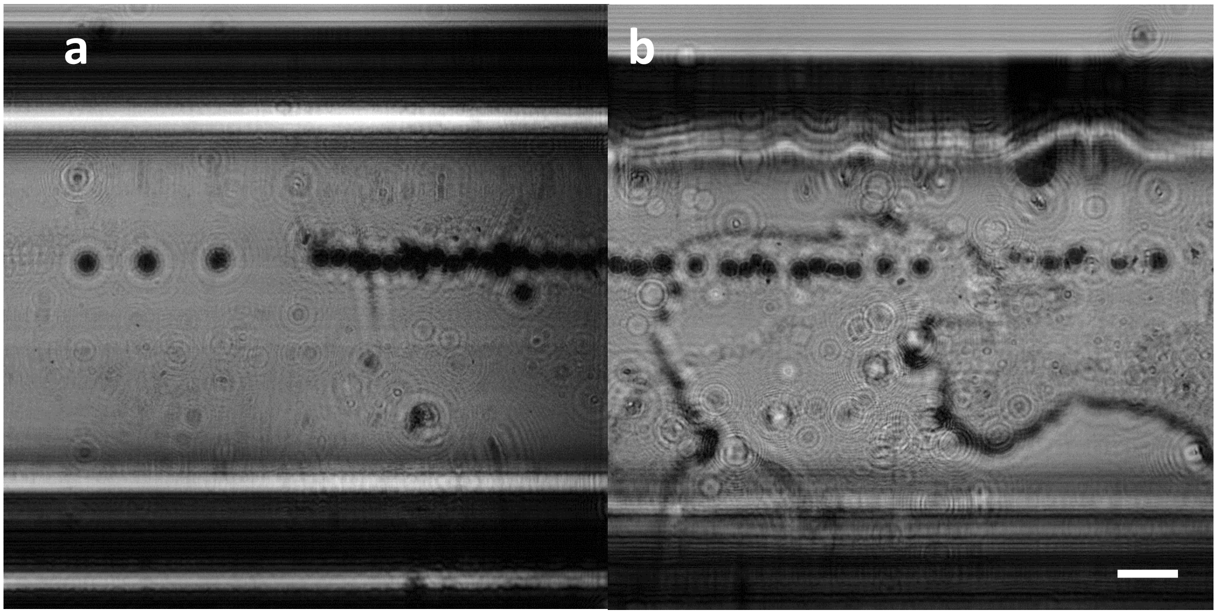


**Figure S5.** Levitation height change of microspheres with an average density of 1.05 g/mL in an incubator platform for 24 h. Acquired and reconstructed holograms at (a) 0 and (b) 24$^{th}$ hour. The mean levitation height of microspheres is 393.8 µm and 397.9 µm at 0 and 24$^{th}$ hour, respectively. Scale bar: 200 µm.

**Table S1.** The prices of the components used for building the platform

| Component | Price ($) |
|---|---|
| Thor Labs LED528EHP | 3.7 |
| Pinhole (100 µm) | 2 |
| 3D printed enclosure (filament) | 2 |
| 2 x NdFeB Block Magnet | 0.5 |
| Microcapillary glass channel (consumable) | 0.7 |
| Sony IMX219 CMOS imaging sensor | 40 |
| Raspberry Pi 3 Model B+ | 45 |
| Total | 93.9 |

**Video 1**. The video of hologram (left) and back-propagated amplitude image (right). The amplitude image was back-propagated along z-axis to identify in-focus image. The levitation heights of microparticles were computed with respect to the bottom magnet. The borders of top and bottom magnets were indicated as pink lines and interior contours of microcapillary channel were indicated as green lines.
